# NeuroMDAVIS: Visualization of single-cell multi-omics data under deep learning framework

**DOI:** 10.1101/2024.02.17.580541

**Authors:** Chayan Maitra, Dibyendu B. Seal, Vivek Das, Rajat K. De

## Abstract

Single-cell technologies have favoured extensive advancements in cell-type discovery, cell state identification, development of lineage tracing and disease understanding among others. Further, single-cell multi-omics data generated using modern technologies provide several views of omics contribution for the same set of cells. However, dimension reduction and visualization of biological datasets (single or multi-omics) remain a challenging task since obtaining a low-dimensional embedding that preserves information about local and global structures in data, is difficult. Further, combining different views obtained from each omics layer to interpret the underlying biology is even more challenging. Earlier, we have developed NeuroDAVIS which can perform the task of visualization of high-dimensional datasets of a single modality while preserving cluster-structures within the data. Nevertheless, there is no model so far that supports joint visualization of multi-omics datasets. Joint visualization refers to transforming the feature space of each individual modality and combining them to produce a latent embedding that supports visualization of the multi-modal dataset in the newly transformed feature space. In this work, we introduce NeuroMDAVIS which is a generalized version of NeuroDAVIS for visualization of biological datasets having multiple modalities. To the best of our knowledge, NeuroMDAVIS is the first of its kind multi-modal data visualization model. It is able to learn both local and global relationships in the data while generating a low-dimensional embedding useful for downstream tasks. NeuroMDAVIS competes against state-of-the-art visualization models like t-Distributed Stochastic Neighbor Embedding (t-SNE), Uniform Manifold Approximation and Projection (UMAP), Fast interpolation-based t-SNE (Fit-SNE), and the Siamese network-based visualization method (IVIS).

## 1 Introduction

With technological advancements, the realm of molecular biology has advanced into a plethora of possibilities. Single-cell technology has opened up newer dimensions for omics data analyses that lead to unravelling of the disease or developmental processes at the cellular level. Single-cell RNA-sequencing (scRNA-seq) allows measurement of mRNA expressions at a cellular resolution. Current scRNA-seq protocols including DROP-seq [1], SMART-seq2 [2], 10x Genomics, 10x Chromium, CEL-seq2 [3] and MARS-seq [4]. ATAC-seq [5], on the other hand, allows sequencing of open chromatin regions within the genome. More recently, joint sequencing technologies have come up, which allow simultaneous measurement of more than one modalities thereby offering multiple views of the same cells of interest. These technologies include CITE-seq [6], REAP-seq [7] (joint measurement of proteins and transcriptome), scM&T-seq [8], scMT-seq [9], scTrio-seq [10] (simultaneous measurement of methylome and trascriptome), and sci-CAR [11], SNARE-seq [12] and SHARE-seq [13] (joint measurement of chromatin accessibility and transcriptome in single-cells or nuclei). All these technologies facilitate deeper understanding of the cellular identity and processes in organs and tissues.

Multi-omics datasets originate from different sources, and a large number of dimensions in each modality makes the data incredibly complex. Thus, reduction of dimension while preserving the inherent structures within the data, is a basic requirement for easy exploration and interpretation of data. Visualizations help in identifying patterns, clusters, outliers, and correlations within these single and multi-omics datasets.

The task of visualization can be compared to a refined Dimension Reduction (DR) task, where the number of reduced dimensions is limited to two or three. Some of the DR techniques developed so far are used to accomplish visualization task. Earlier, mostly the methods were linear in nature [14]. However, a large number of nonlinear DR techniques for data visualization have come up during the last decade. These methods possess the ability to handle data non-linearity. Methods like SNE [15], t-SNE [16] and Fit-SNE [17] assume that the data follow a particular theoretical distribution. Hence, they try to learn a low-dimensional embedding that follows the similar distribution, which, however, may contradict the assumption. Some methods, like Isomap [18] and UMAP [19], try to reconstruct the hidden topological structure within the data but fail to do so when the actual structure is complex. Other neural network-based methods like IVIS [20], SOM [21], Autoencoders [22] try to learn a suitable non-linear transformation to project the data into low dimension. Neural network-based methods have an edge over the others since they do not assume any distribution of the data, and they are parametric in the sense that several parameters are involved in architecture and learning. For this reason, neural network-based models can be trained on one dataset and applied on some other datasets originated from the same process.

Motivated by the wide benefits of using neural networks, in our earlier work, we had developed a data visualization method using a neural network model, called NeuroDAVIS [23]. NeuroDAVIS had been benchmarked against the state-of-the-art methods in multiple ways but it falls short to visualize multi-modal datasets. When it comes to visualizing observations from multi-omics experiments, there is no joint data visualization method that can combine views from several omics layers. Such a joint visualization can assist researchers to explain the underlying biology of a system/process in a better way. For this reason, in this work, we have introduced a multi-modal data visualization model developed under a deep learning framework that is capable of extracting crucial features from the data and produce an effective joint visualization. NeuroMDAVIS is a non-recurrent, feed-forward deep learning model which does not assume any kind of data distribution to visualize a high-dimensional data. Most of the state-of-the-art methods have been observed to perform well on complex datasets when conjugated together with an initialization step like that of applying a Principal Component Analysis (PCA), to reduce the number of dimensions to something less than or equal to 50 [24]. NeuroMDAVIS does not require any such initialization, which makes it more efficient and effective. Our primary aim being multi-omics data visualization, in this study, we have limited our experiments to multi-omics datasets only. However, NeuroMDAVIS can be applied to other domains as well. Additionally, we have also explored how NeuroDAVIS [23] performs on single-omics data.

## 2 Methods

The deep learning model, called NeuroMDAVIS, developed in this work, allows joint visualization of high-dimensional multi-omics datasets. It is a generalized version of NeuroDAVIS [23] introduced earlier to provide visualization of a single data modality. Nevertheless, NeuroMDAVIS incorporates significant changes to the network architecture over NeuroDAVIS.

### 2.1 Architecture

NeuroMDAVIS architecture represents a novel structure as shown in Figure 1. It consists of four different types of layers, viz., an *Input layer*, a *Latent layer*, one or more *Hidden layer(s)* and a *Reconstruction layer*. Hidden layer(s) (the dashed rectangle in Figure 1) are of two types, viz., *Shared hidden layer(s)* and *Modality-specific hidden layer(s). Input layer, Latent layer* and *Shared hidden layer(s)* are densely connected, i.e., each node in any of these layers are connected to each node of its adjacent layer(s). However, *Modality-specific hidden layer(s)* does not share a completely dense connection, instead, it shares a dense connection modality-wise only.

**Figure 1:**
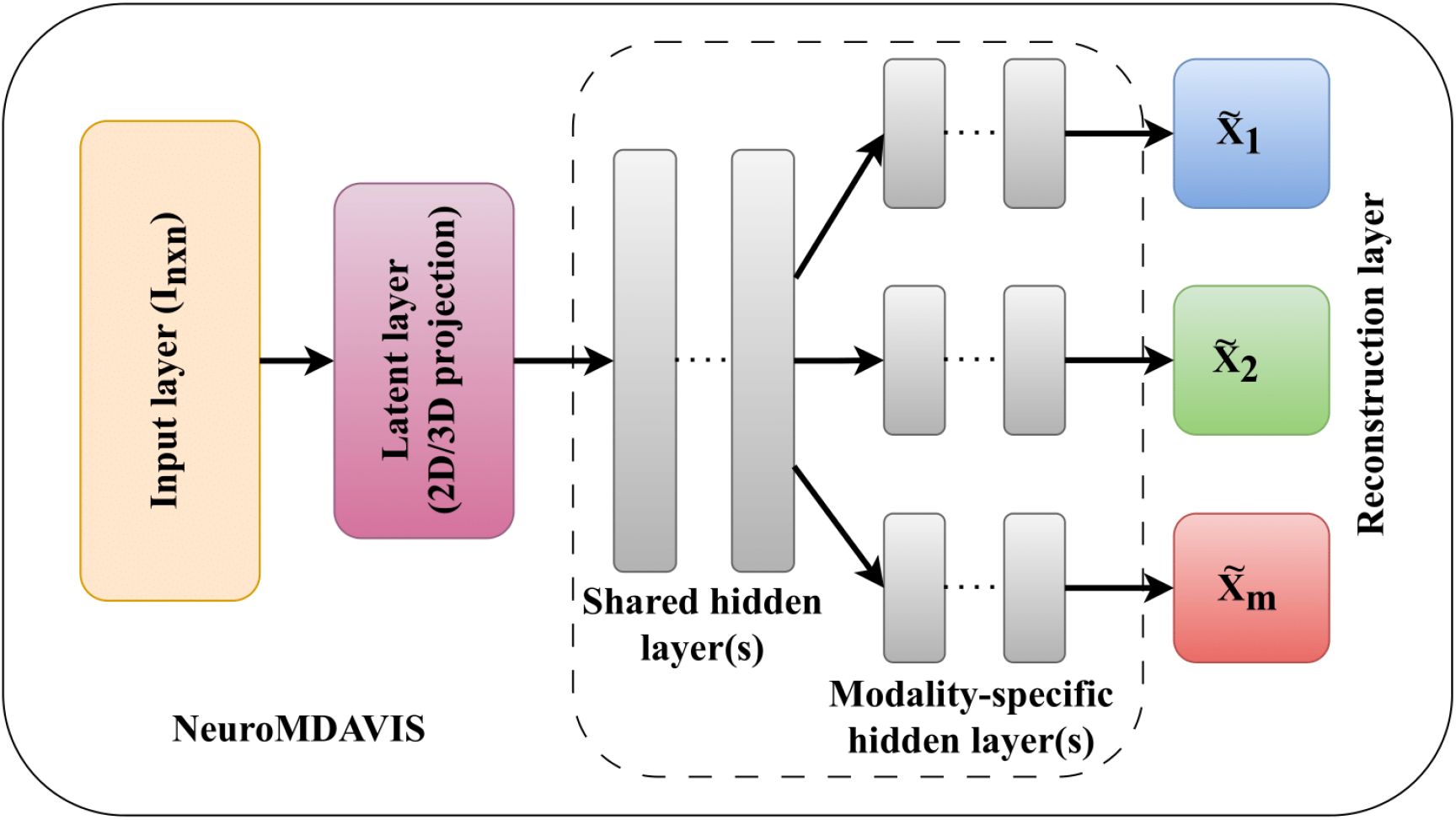
The NeuroMDAVIS network architecture developed for visualization of high-dimensional multi-omics datasets.

Let 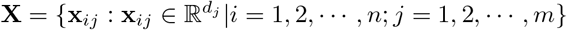 be a dataset containing *m* different omics modalities. In each modality, there are equal number (say *n*) of samples/observations, where each sample has data related to all *m* omics modalities. A *j*^*th*^ (*j* = 1, 2, …, *m*) modality is characterized by *d*_*j*_ features, i.e., **x**_*ij*_ = [*x*_*ij*1_, *x*_*ij*2_, …, *x*_*ijd*_]^*T*^, *i* = 1, 2, …, *n*; *j* = 1, 2, …, *m*. The *Input layer* of NeuroMDAVIS has *n* number of neurons while the *Latent layer* has *k* (= 2 or 3) number of neurons, where *k* is the desired number of dimensions to be used for visualization. Both the *Modality-specific hidden layer(s)* and *Reconstruction layer* have *m* modality-specific sub-modules. In each sub-module of the *Reconstruction layer*, there are *d*_*j*_ neurons. However, the number of neurons in the *Hidden layer(s)* are decided empirically.

*Input layer* of NeuroMDAVIS takes an identity matrix of order *n × n* as input. Similar to NeuroDAVIS, the *Input layer* creates a random latent embedding of *n* samples at the *Latent layer. Latent layer* then tries to regress all the data modalities simultaneously through the aforesaid *Hidden layers*. The purpose of using a *Shared hidden layer* is to capture the common information across different modalities for a particular sample/observation, whereas, a *Modality-specific hidden layer* extracts the modality-specific information for that sample/observation. In this work, only one *Shared hidden layer* and one *Modality-specific hidden layer* each has been used. One can use multiple such layers based on the problem requirement. When *m* = 1, the architecture resembles that of NeuroDAVIS with a general *Hidden layer(s)* instead of distinct *Shared hidden layer(s)* and *Modality-specific hidden layer(s)*. Finally, the *Reconstruction layer* reconstructs the data in hand, i.e., the original individual omics modalities.

### 2.2 Forward propagation

We have considered a dataset **X** with *m* omics modalities each having *n* paired samples, i.e.,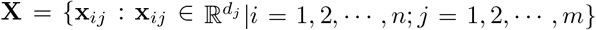. NeuroMDAVIS takes an identity matrix **I** of order *n × n* as input. An *i*^*th*^ column vector **e**_*i*_ of **I** is considered for regressing the *i*^*th*^ paired sample. More precisely, **e**_*i*_ propagates through the layers of NeuroMDAVIS and reconstructs an approximate version 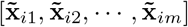 of [**x**_*i*1_, **x**_*i*2_, …, **x**_*im*_] at the *Reconstruction layer* on presentation of *i*^*th*^ sample. Let **a**_*il*_ and **h**_*il*_ correspond to the input to and output from the *l*^*th*^ layer respectively. Let **W**_*l*_ and **b**_*l*_ be the weight matrix between (*l* − 1)^*th*^ layer and *l*^*th*^ layer (*l* = 1, 2, …, (*s* + 1)), and the bias term for nodes in *l*^*th*^ layer respectively. Here *s* stands for the total number of *Shared hidden layer* present in the network. An *l*^*th*^ layer may be any of the *Input layer* (*l* = 0), *Latent layer* (*l* = 1) or *Shared hidden layer(s)* (*l* = 2, 3, …, (*s* + 1)). Thus, for the *Input layer*, we have

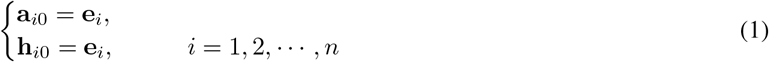

For the *Latent layer*, we have

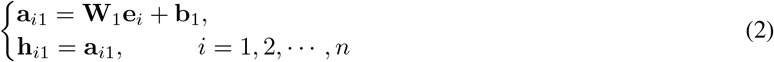

Here, the weight parameters, controlled by the input **e**_*i*_, create an independent low-dimensional representation of the *i*^*th*^ sample at the *Latent layer*. This is required to ensure that only the links connected to the *i*^*th*^ neuron of the *Input layer* activate neurons in the *Latent layer* on presentation of the *i*^*th*^ sample. Then, for *s Shared hidden layer(s)*, we have

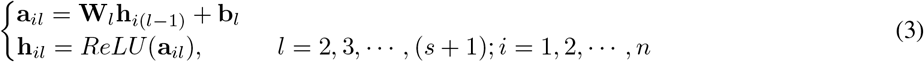

where *ReLU* (**y**) = *max*(**0, y**); *max* (maximum) being an element-wise operation. The output from the last *Shared hidden layer* will flow till the end of the network through different *Modality-specific hidden layer(s)* and reconstruct every modality at the *Reconstruction layer*. Let the number of *Modality-specific hidden layer(s)* is *p*. Let us assume that 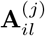 and 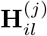 denote the input to and output from the *j*^*th*^ module of the *l*^*th*^ layer respectively. Let us also assume that 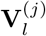 and 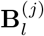 denote the weight matrix between the *j*^*th*^ modules of the (*l* − 1)^*th*^ and *l*^*th*^ layer, and the bias term for nodes in *j*^*th*^ module of the *l*^*th*^ layer respectively. Here, an *l*^*th*^ layer may be any of the *Modality-specific hidden layer(s)* (*l* = (*s* + 2), (*s* + 3), …, (*s* + *p* + 1)) or a *Reconstruction layer* (*l* = (*s* + *p* + 2)).

Thus, for the first *Modality-specific hidden layer*, we have

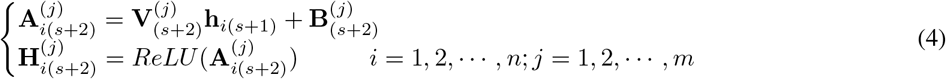

For the remaining *Modality-specific hidden layer(s)*, we have

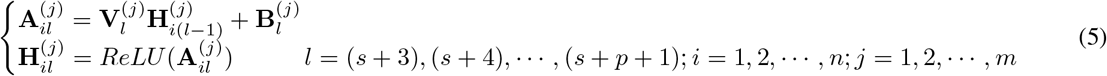

Finally, at the *Reconstruction layer*, a reconstruction (lossy) of the original data is formed. For the *Reconstruction layer*, we have

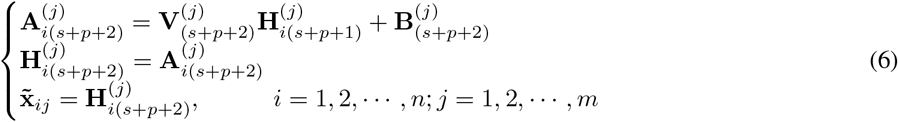

Thus, NeuroMDAVIS projects the latent embedding for a sample, obtained at the *Latent layer*, to different *d*_*j*_ dimensional spaces, corresponding to each of *j*^*th*^ omics modality, through the *Hidden layer(s)*, so that the sample gets reconstructed at the *Reconstruction layer*. The vector 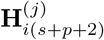 represents the lossy reconstruction 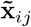 of **x**_*ij*_, the *i*^*th*^ sample in the *j*^*th*^ omics modality.

### 2.3 Learning

NeuroMDAVIS enables dimension reduction and visualization of high-dimensional multi-omics data. Similar to NeuroDAVIS, NeuroMDAVIS also tries to reconstruct the data to be visualized. For *i*^*th*^ sample **x**_*ij*_, NeuroMDAVIS tries to minimize the reconstruction error 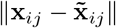 in order to find an optimal reconstruction 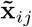 of the sample **x**_*ij*_. Since different omics modalities may have different number of dimensions, to avoid disparity in learning, a balancing parameter *λ*_*j*_ has been introduced into the objective function for NeuroMDAVIS. The value of *λ*_*j*_ lies in (0, 1], where higher value of *λ*_*j*_ implies higher weightage to the *j*^*th*^ data modality. *λ*_*j*_ is useful when knowledge about the data modalities are available *a priori*. In absence of prior knowledge, one can simply choose *λ*_*j*_ = 1, ∀*j*. The objective function thus becomes

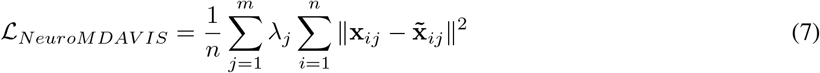

To avoid overfitting and minimization of model complexity, *L*2 regularization, involving activities of nodes and weights, has been considered. The objective function thus becomes

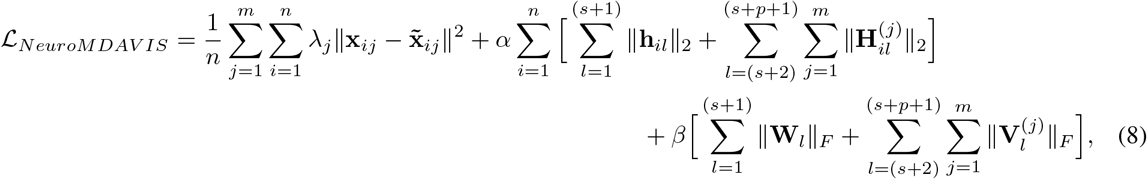

where *α* and *β* are regularization parameters set empirically. We have used Adam optimizer for training NeuroMDAVIS. The number of epochs needed for convergence has also been set empirically. After convergence, NeuroMDAVIS learns a parametric function that can efficiently produce the omics modalities being reconstructed, from the low-dimensional latent embedding. This latent embedding is then extracted to produce a joint visualization of the multi-omics data.

### 2.4 Projection of new observations

The first two layers of NeuroMDAVIS, viz., *Input layer* and *Latent layer*, are used to learn suitable regressors. In other words, they control the points in the low-dimensional embedding in each iteration, while the remaining part of the network tries to optimize a function that can project the low-dimensional data into high dimension. Once NeuroMDAVIS is trained on a training dataset, we can visualize the data at the *Latent layer*. The weights from the *Latent layer* through the *Reconstruction layer* have been learned in such a way that it can project any low-dimensional point to a high-dimensional space.

In order to visualize new observations (not present during training but having a similar distribution), the *Input layer* of NeuroMDAVIS has to be presented with an identity matrix which will be of the order of the number of observations in this unseen data (test dataset). This test dataset must contain the same number of omics modalities as the training data. Thus, the *Input layer* dimension will be equal to the size of the test dataset. The weights between the *Input layer* and the *Latent layer* need to be initialized again, and only the sub-network comprising the *Input layer* and the *Latent layer* needs to be re-trained, allowing the weights connecting these two layers to be updated, while the others are kept frozen to the already learned values. These weights are then used to visualize new samples.

Let there be *n*_1_ observations in the test data. In order to visualize the test data, an identity matrix of size *n*_1_ *× n*_1_ is fed to the *Input layer* of the new network. Similar to the training phase, a loss is calculated at the *Reconstruction layer*. However, unlike training, during visualization of new samples, only the weights connecting the *Input layer* to the *Latent layer* are updated via standard back-propagation. Upon convergence, the final latent embedding for these unseen samples can be extracted from the *Latent layer*.

## 3 Results

Performance of NeuroMDAVIS has been demonstrated on multi-omics (CITE-seq and multiome) datasets (Table 1) for their visualization. Additionally, we have also demonstrated the performance of NeuroDAVIS [23] on single-omics datasets (Table 1), like single-cell RNA-seq and Mass Cytometry data. Comparative performance of NeuroMDAVIS with some state-of-the-art visualization methods, like t-SNE, UMAP, IVIS and Fit-SNE, has been analyzed. Table 2 presents a theoretical comparison of NeuroDAVIS and NeuroMDAVIS with these existing methods. At the begining of this section, we have presented results of the experiments performed on single-omics datasets using NeuroDAVIS, which include those carried out on single-cell RNA-seq data (Section 3.1) and the ones carried out on Mass Cytometry data (Section 3.2). This is followed by the results on multi-omics datasets (Section 3.3) including those on CITE-seq (Section 3.3.1) and mutliome data (Section 3.3.2).

**Table 1:** Summary of datasets used in this work.

|  | Dataset | Technology | Description | #Cells | #RNAs | #ADTs/<br>#Peaks | Batches<br>present | Source |
| --- | --- | --- | --- | --- | --- | --- | --- | --- |
| Single-omics modality | *Usoskin* | scRNA-seq | scRNA-seq dataset contains mouse neuronal cells in the dorsal root ganglion | 622 | 25334 | - | No | [25] |
|  | *Jurkat* | scRNA-seq | RNA-seq derived from T lymphocyte cell line originally obtained from the peripheral blood of a 14-year-old boy with acute lymphoblastic leukemia | 3388 | 32738 | - | No | [26] |
|  | *Wong* | Mass Cytometry | Mass Cytometry dataset containing protein targets from 8 distinct human tissues | 327457 | - | 39 | No | [27] |
| Multiple omics modalities | *bmcite30k* | CITE-seq | scRNA-seq profiles measured alongside a panel of antibodies from bone marrow | 30672 | 17009 | 25 | Yes | [28] |
|  | *kotliarov50k* | CITE-seq | CITE-seq profiling of 82 surface proteins and transcriptomes of 53,201 single-cells from healthy high and low influenza-vaccination responders | 58654 | 32738 | 87 | Yes | [29, 30] |
|  | *pbmc10k* | Multiome | single-cell multiome ATAC and gene expression data from cryopreserved human peripheral blood mononuclear cells (PBMCs) of a healthy female donor | 11909 | 36601 | 108377 | No | [31] |

**Table 2:** A theoretical comparison among the visualization methods used in this study.

| Method | Type | Supports multi-modal<br>visualization (Yes/No) | Makes assumption about<br>data distribution (Yes/No) | Parametric/<br>Non-parametric |
| --- | --- | --- | --- | --- |
| NeuroMDAVIS | Neural network-based | Yes | No | Parametric |
| NeuroDAVIS | Neural network-based | No | No | Parametric |
| t-SNE | Distribution-based | No | Yes | Non-parametric |
| UMAP | Manifold-based | No | No | Non-parametric |
| Fit-SNE | Distribution-based | No | Yes | Non-parametric |
| IVIS | Neural network-based | No | No | Parametric |

### 3.1 Visualizing single-cell RNA-seq data using NeuroDAVIS

To analyze the performance of NeuroDAVIS for visualization of single-omics, we have used two single-cell RNA-seq datasets, viz., *Usoskin* and *Jurkat*. In our earlier investigaion [23], we had shown how NeuroDAVIS embeddings for these two datasets compare against those obtained using t-SNE-, UMAP-, IVIS– and Fit-SNE-generated embeddings. We had also explained how NeuroDAVIS has outperformed the other methods for *Jurkat* dataset and competes against them for *Usoskin* dataset with respect to the correlation coefficient values between the pairwise distances in the original space and the embedding generated by these methods including NeuroDAVIS itself. Further, we had shown that performance of *k*-means and hierarchical clustering on NeuroDAVIS-generated embeddings of these two datasets, compares favourably with that obtained on embeddings generated by the other visualization methods.

In order to validate the effectiveness of NeuroDAVIS for visualization of these two datasets –*Usoskin* and *Jurkat*, in this work, we have performed a few additional experiments to quantify attributes, like shape preservation and scalability, as explained in the following sections.

#### 3.1.1 Shape preservation

An important attribute to compare dimension reduction methods, is shape preservation. While every method tries to preserve local and global structures/shapes within the data, they end up preserving one over the other. In order to analyze how NeuroDAVIS compares with the other state-of-the-art methods to preserve local and global shapes, we have compared the pairwise distances among all points in high-dimension to that in their reduced low-dimensional embedding, generated by these methods.

First, we have calculated all possible distances in the original (high-dimensional) space, and distributed them across 50 equal-width bins in sorted order. Similarly, the pairwise distances in the low-dimensional embedding have been recorded and mapped into their corresponding bins. To understand how good global distances are preserved by each embedding, we have used boxplots for displaying the distribution of the pair-wise distances in low dimension over that in high dimension, throughout the bins, for all the methods, as shown in Figures 2(A) and 2(B). The boxplots, at the beginning, represent short distances, while those at the end represent long distances. Thus, lower the spread of the boxplot, better is the preservation of distances. Moreover, the median of the boxplots should follow an increasing order, i.e., the median for trailing boxplots representing long distances should be higher than that for the leading boxplots representing short distances. This signifies that the order of pairwise distances between observations in high dimension is similar to that between observations in low dimension.

**Figure 2:**
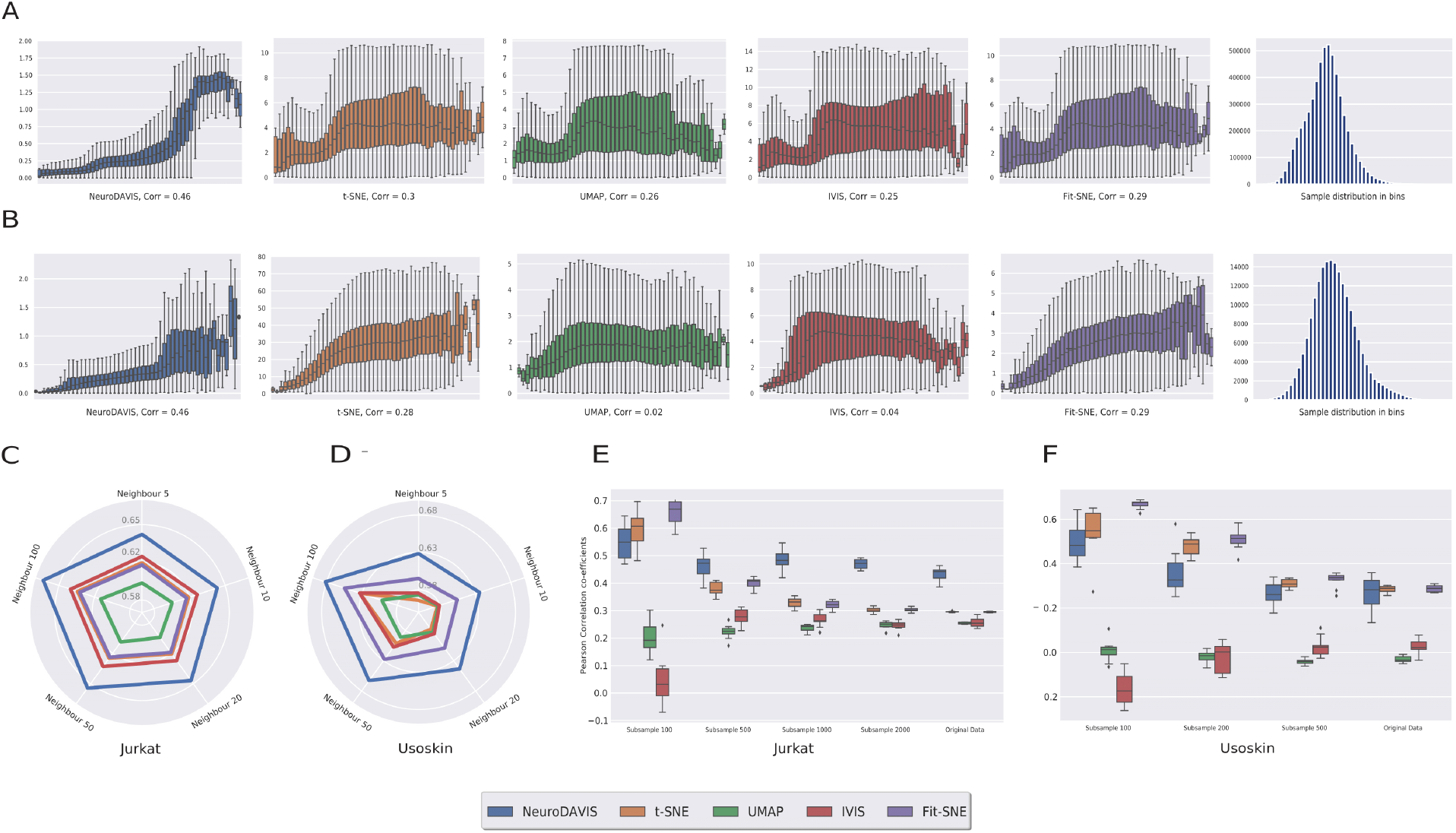
(A) and (B) show the results of performance comparison of NeuroDAVIS with that of t-SNE, UMAP, IVIS and Fit-SNE in terms of distance preservation capability and correlation between pairwise distances in high dimension to that in low dimension of *Jurkat* and *Usoskin* datasets respectively. Distances in high dimension have been measured and distributed across 50 equal-width bins in sorted order. The corresponding distances in low dimension have also been mapped into the same bins. The leading boxplots represent short distances while the trailing ones represent long distances. Lower the spread of the boxplot, better is the preservation of distances. The median of the trailing boxplots representing long distances should be higher than that of the leading boxplots representing short distances. It signifies that the order of pairwise distances among observations in high dimension is similar to that among observations in low dimension. The last histogram for each dataset represents the count of distances in each bin. (C) and (D) show results of comparison of local shape preservation capability of NeuroDAVIS with that of t-SNE, UMAP, IVIS and Fit-SNE in terms of trustworthiness score on *Jurkat* and *Usoskin* datasets respectively. (E) and (F) show results of comparison of robustness of NeuroDAVIS with that of t-SNE, UMAP, IVIS and Fit-SNE in terms of subsampling-based correlation analysis on *Jurkat* and *Usoskin* datasets respectively.

It has been observed that for both *Usoskin* and *Jurkat*, NeuroDAVIS has preserved short distances better since the spread in the leading boxplots tends to be zero. On observing the median distances, we have found that for both the datasets, NeuroDAVIS exhibits an increasing pattern, outperforming all the other methods which show a descent in median distances as we go towards the trailing boxplots. The Pearson correlation coefficient values between all-pair distances in high-dimension and that in low dimension have been found to be 0.46 for both the datasets, which is higher than that obtained using the other methods. The last subfigure within Figures 2(A) and 2(B) each represent the support values, i.e., the number of pairwise distances within each bin.

Additionally, we have compared the NeuroDAVIS-generated embedding with that produced by the other methods with respect to their trustworthiness [32], a measure to quantify local neighbourhood preservation post-dimension reduction. For five different values of neighbours, the trustworthiness score has been calculated for both *Jurkat* and *Usoskin* datasets. The trustworthiness values, as shown in Figures 2(C) and 2(D), reflect that local neighbours are preserved better in the NeuroDAVIS embedding as compared to the embeddings produced by the other methods.

#### 3.1.2 Subsampling and robustness

In order to test robustness of the methods, we have carried out a subsample-based experiment on *Usoskin* and *Jurkat* datasets. For each of these datasets, we have initially drawn a small subsample (100 samples) randomly and applied all these methods for dimension reduction. We have recorded the correlation coefficient values (Pearson) between pairwise distances in the original distribution and that in the low-dimensional embedding produced by each of these methods. This experiment has then been repeated multiple times with increasing subsample sizes on both the datasets. Interestingly, we have observed that the performance of NeuroDAVIS has been stable and consistent as and when we increase the subsample size (Figures 2(E) and 2(F)). However, though t-SNE and Fit-SNE have performed well on small subsample sizes, they have failed to scale for large subsample sizes. The performance of UMAP and IVIS, on the other hand, has not been good enough even for small subsample sizes. NeuroDAVIS has outperformed all the other methods for both these datasets when the subsample size reaches the maximum number of samples in the datasets. Thus, NeuroDAVIS stands out as the most robust method among all the visualization methods used in the experiment.

### 3.2 Visualizing Mass Cytometry data using NeuroDAVIS

To further demonstrate the effectiveness of NeuroDAVIS, we have considered a Mass Cytometry data (*Wong* data), having more than 300K samples with 39 protein targets. The labels for each cell generated using Louvain clustering-based phenograph and manual annotation, have been collected from [19]. Due to computational limitation, in this study, we have used a random sample of size 30k from this data for visualization using NeuroDAVIS. A similar embedding has also been produced by the other state-of-the-art methods. Comparable visualization traits have been found in the embeddings generated by all the methods (Figure 3(A)), where the *Tgd* (*γδT*) cell cluster and the *Contaminant* (including *B*) cell clusters have been splitted into distinct clusters. However, unlike t-SNE, UMAP and Fit-SNE, NeuroDAVIS has been able to keep the *CD*8*T* cell cluster intact, which is only comparable to IVIS. Similar observation can be made for the *CD*4*T* cell cluster. Thus, NeuroDAVIS undoubtedly preserves local neighbours better than the other methods. This property has been proved mathematically during our earlier investigation on NeuroDAVIS [23].

**Figure 3:**
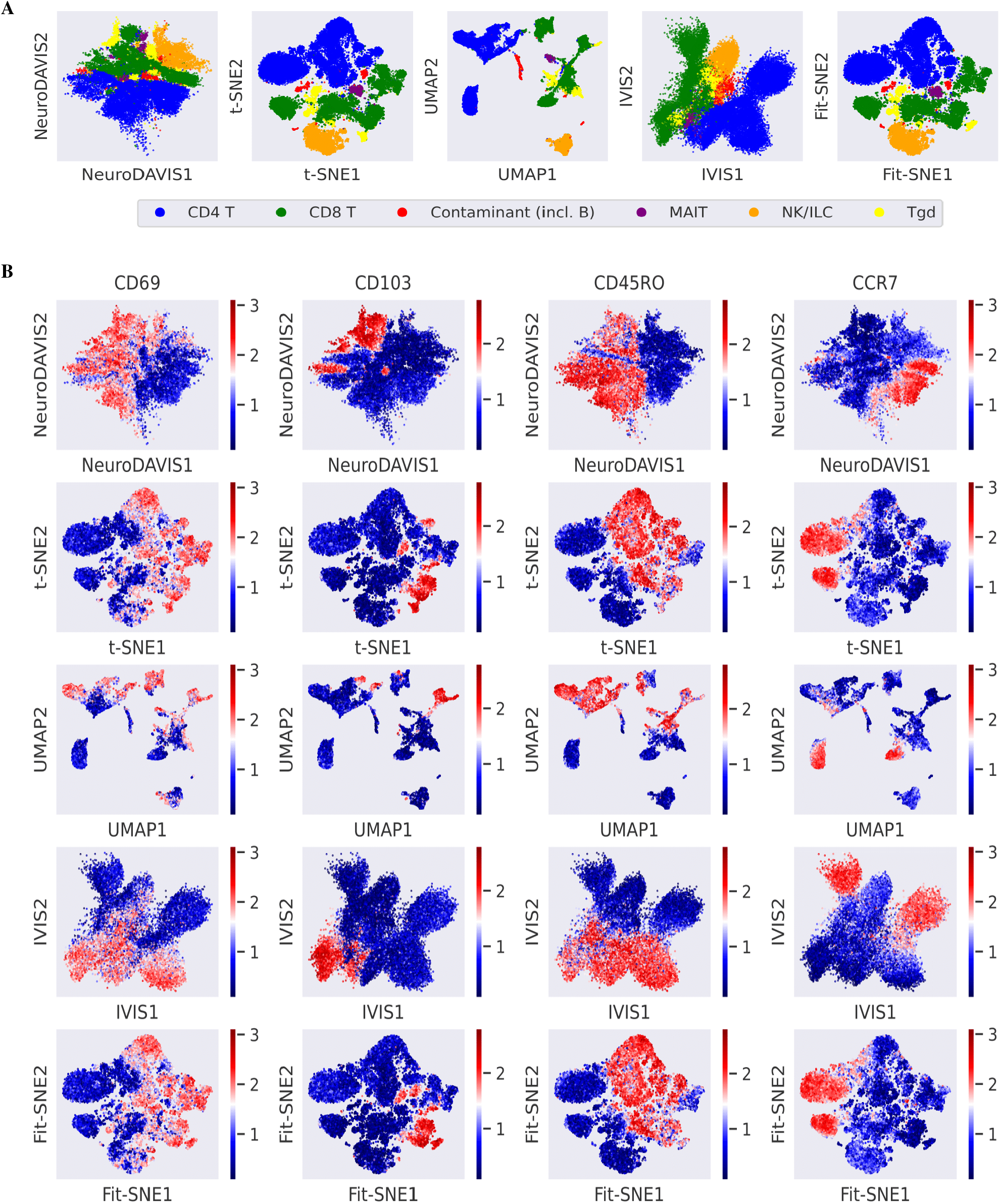
(A) 2-dimensional embeddings produced by NeuroDAVIS, t-SNE, UMAP, IVIS and Fit-SNE for the Mass Cytometry *Wong* data, coloured by cell-type; *MAIT* denotes mucosal-associated invariant *T* cells; *ILC* stands for innate lymphoid cells; and Tgd represents *γδT* cells. (B) Expression of *T* cell marker proteins *CD*69, *CD*103, *CD*45*RO, CCR*7 on the NeuroDAVIS, t-SNE, UMAP, IVIS and Fit-SNE-generated embeddings.

Subsequently, the quality of embeddings has been evaluated by projecting the expression profiles of a few chosen proteins on the embeddings. As the major cell-type present in the data is *T* cell, we have tried to visualize the expression levels of different *T* cell markers, viz., *CD*69, *CD*103 (a resident memory *T* cell marker), *CD*45*RO* (a memory *T* cell marker) and *CCR*7 (naive *T* cell marker) [19]. As illustrated in Figure 3(B), the expression levels for all these marker proteins have revealed that NeuroDAVIS has been able to accurately reproduce the *T* cell differentiation stage (blue indicating low expression and red indicating high expression) inside each main cluster. While all the embeddings have represented an analogous continua of clusters, they do not appear to be organized along a single, recognizable axis like that in NeuroDAVIS.

### 3.3 Visualizing multi-omics data using NeuroMDAVIS

To evaluate the performance of NeuroMDAVIS with respect to multi-omics data visualization, we have used two multi-omics datasets including two CITE-seq (joint profiling of RNA and surface protein measurements) data and one multiome (paired RNA-seq and ATAC-seq) data. Since the existing visualization methods used for comparison do not support multi-modal visualization, for each dataset, we have concatenated the omics modalities available against the set of paired cells and used them as input to these methods for visualization. The following subsections describe the experiments carried out on these two datasets and their corresponding results.

#### 3.3.1 Visualizing CITE-seq data

First, we have considered two CITE-seq datasets, viz., *bmcite*30*k* and *kotliarov*50*k*. These datasets have been downloaded from and preprocessed following [33]. CITE-Seq allows concurrent measurement of mRNA and cell surface proteins. In order to trace cell-types, most of the existing visualization methods either use separate projections for ADTs and RNA molecules, or project the protein expressions on the transcriptomic landscape. However, NeuroMDAVIS supports joint visualization of multiple modalities. As shown in Figures 4(A) 5(A), for these two datasets, NeuroMDAVIS has been able to keep distinct cell types into distinct clusters, the clustering quality being qualitatively comparable only to IVIS. However, t-SNE, UMAP and Fit-SNE have been unable to keep clusters well separated. The reason for this might be the absence of a prior initialization step, like the usage of PCA, which serves as a good initialization for other state-of-the-art methods, like t-SNE or UMAP [34]. On the contrary, NeuroMDAVIS and IVIS, both being neural network based methods, have been able to produce better visualization even when used solely on the multi-modal data without any pre-processing.

**Figure 4:**
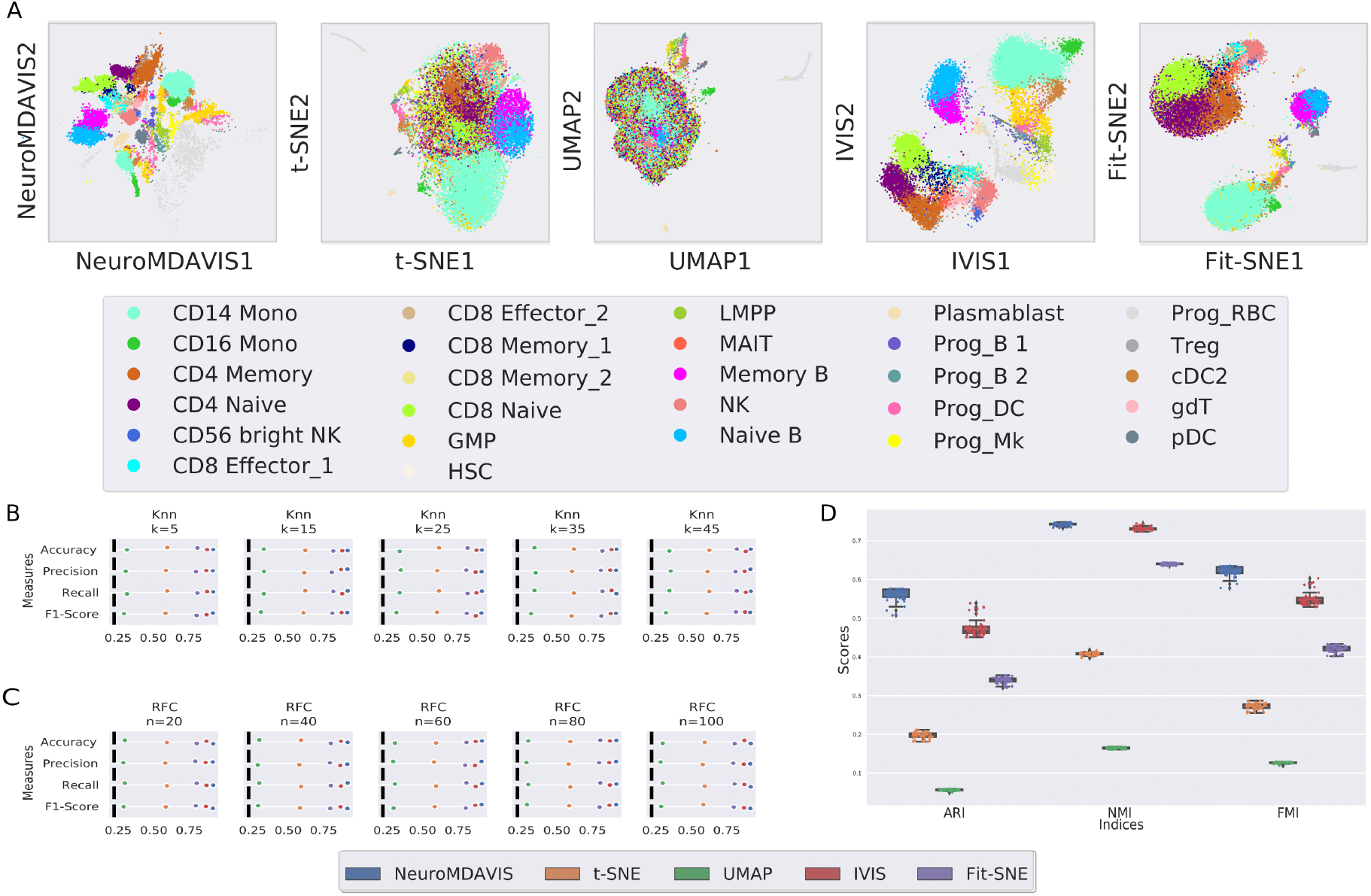
(A) 2-dimensional embeddings produced by NeuroMDAVIS, t-SNE, UMAP, IVIS and Fit-SNE for *bmcite*30*k* dataset. (B) and (C) show Classification performance on low-dimensional embeddings of *bmcite*30*k* dataset generated using NeuroMDAVIS, t-SNE, UMAP, IVIS and Fit-SNE, in terms of accuracy, precision, recall and F1-score using *k*-NN and Random Forest classifers respectively. (D) Clustering performance on low-dimensional embeddings generated using NeuroMDAVIS, t-SNE, UMAP, IVIS and Fit-SNE with respect to ARI, NMI and FMI scores of *bmcite*30*k* datasets, using *k*-means clustering algorithm.

**Figure 5:**
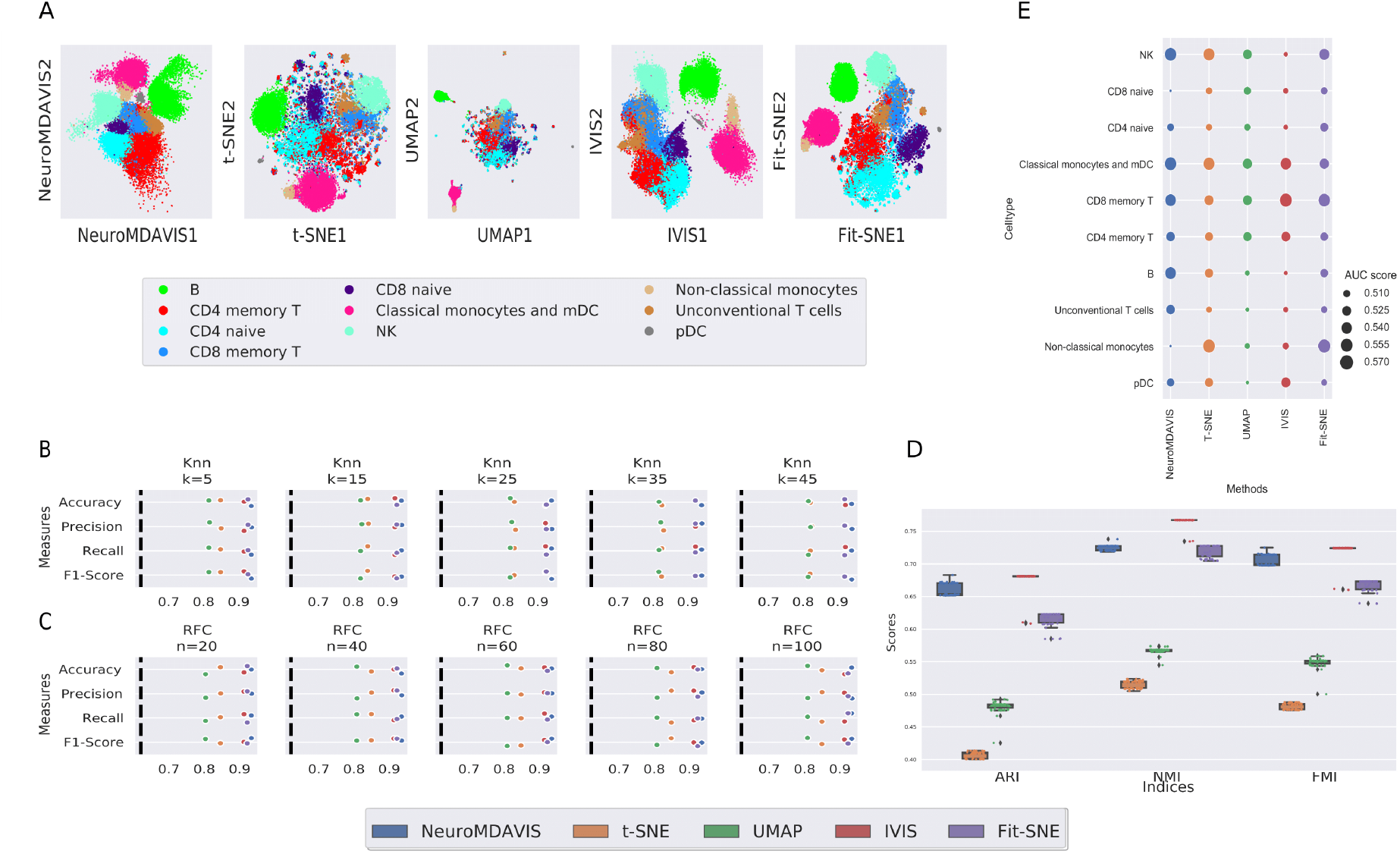
(A) 2-dimensional embeddings produced by NeuroMDAVIS, t-SNE, UMAP, IVIS and Fit-SNE for *kotliarov*50*k* dataset. (B) and (C) show classification performance on low-dimensional embeddings of *kotliarov*50*k* dataset generated using NeuroMDAVIS, t-SNE, UMAP, IVIS and Fit-SNE, in terms of accuracy, precision, recall and F1-score using *k*-NN and Random Forest classifers respectively. (D) Clustering performance on low-dimensional embeddings generated using NeuroMDAVIS, t-SNE, UMAP, IVIS and Fit-SNE with respect to ARI, NMI and FMI scores of *kotliarov*50*k* datasets, using *k*-means clustering algorithm. (E) Area-Under-the-Curve (AUC)-score for high/low influenza vaccine responder classification obtained using *k*-NN classifier (with *k* = 50) on the NeuroMDAVIS, t-SNE, UMAP, IVIS and Fit-SNE projections of *kotliarov*50*k* dataset.

Thereafter, we have used *k*-NN and Random Forest classifiers to identify cell-types based on the NeuroMDAVIS-generated projection, and compare results with that performed on projections generated by other existing methods. Training and test datasets have been prepared in a 80:20 ratio. The parameters *k*, representing the number of neighbours in *k*-NN, and *n*, being the number of estimators in Random Forest classification, have been varied between 5 − 45 and 20 − 100 respectively to ensure consistency of results. As can be seen in Figures 4(B), 4(C), and Figures 5(B), 5(C), NeuroMDAVIS has outperformed all other methods with respect to Accuracy, Precision, Recall and F1-score values for both the datasets, consistently over all values of *k* for *k*-NN and *n* for Random Forest classifiers, on the 20% held out test dataset.

Further, *k*-means clustering on the embedding generated by NeuroMDAVIS, has shown highly competitive performance, if not better than the other existing methods, in terms of Adjusted Rand Index (ARI), Normalized Mutual Information (NMI) and Fowlkes Mallows Index (FMI) scores. As depicted in Figures 4(D) and 5(D), for *bmcite*30*k* dataset, NeuroMDAVIS has outperformed all the other methods while for *kotliarov*50*k* dataset, the performance of NeuroMDAVIS has surpassed t-SNE, UMAP and Fit-SNE by a high margin.

For further validation, we have performed a clinical classification on the *kotliarov*50*k* embedding generated by NeuroMDAVIS. The dataset *kotliarov*50*k* deals with single-cell CITE-seq data gathered from high and low-influenza vaccine responders. We have tried to explore whether NeuroMDAVIS-generated projection is capable of classifying cells into low and high responder categories. For each unique cell cluster, we have performed a train-test split of 80:20 ratio. *k*-NN classifier with *k* = 45 has been used for this binary classification. Figure 5(E) demonstrates that the Area-Under-the-Curve (AUC)-score obtained using NeuroMDAVIS on each of the cell-types has been highly competitive with those obtained using the other state-of-the-art methods, viz., t-SNE, UMAP, IVIS and Fit-SNE. It may be noted here that none of the methods could achieve beyond 58% AUC, which suggests that the cell type labels obtained for this dataset may not be optimal.

#### 3.3.2 Visualizing multiome data

A commonly used method for assessing chromatin accessibility across the genome is the Assay for Transposase-Accessible Chromatin with sequencing (ATAC-seq). One can learn about how chromatin packaging and other variables impact gene expression by using ATAC-seq to sequence open chromatin regions.

In this work, a multiome dataset *pbmc*10*k* containing paired RNA-seq and ATAC-seq data, has been further used to demonstrate the effectiveness of NeuroMDAVIS for multi-omics data visualization. This *pbmc*10*k* dataset has been first preprocessed using MUON [31], reduced to highly variable features only and then further processed to match cells in both the RNA and ATAC modalities, as done in [33]. NeuroMDAVIS has then been applied on the multi-omics data to generate a joint embedding of the paired assays. When compared to t-SNE, UMAP and Fit-SNE, NeuroMDAVIS has produced better visualization of *pbmc*10*k* data mapping each cell-type to a distinct cluster, which is comparable only to that produced by IVIS (Figure 6(A)).

**Figure 6:**
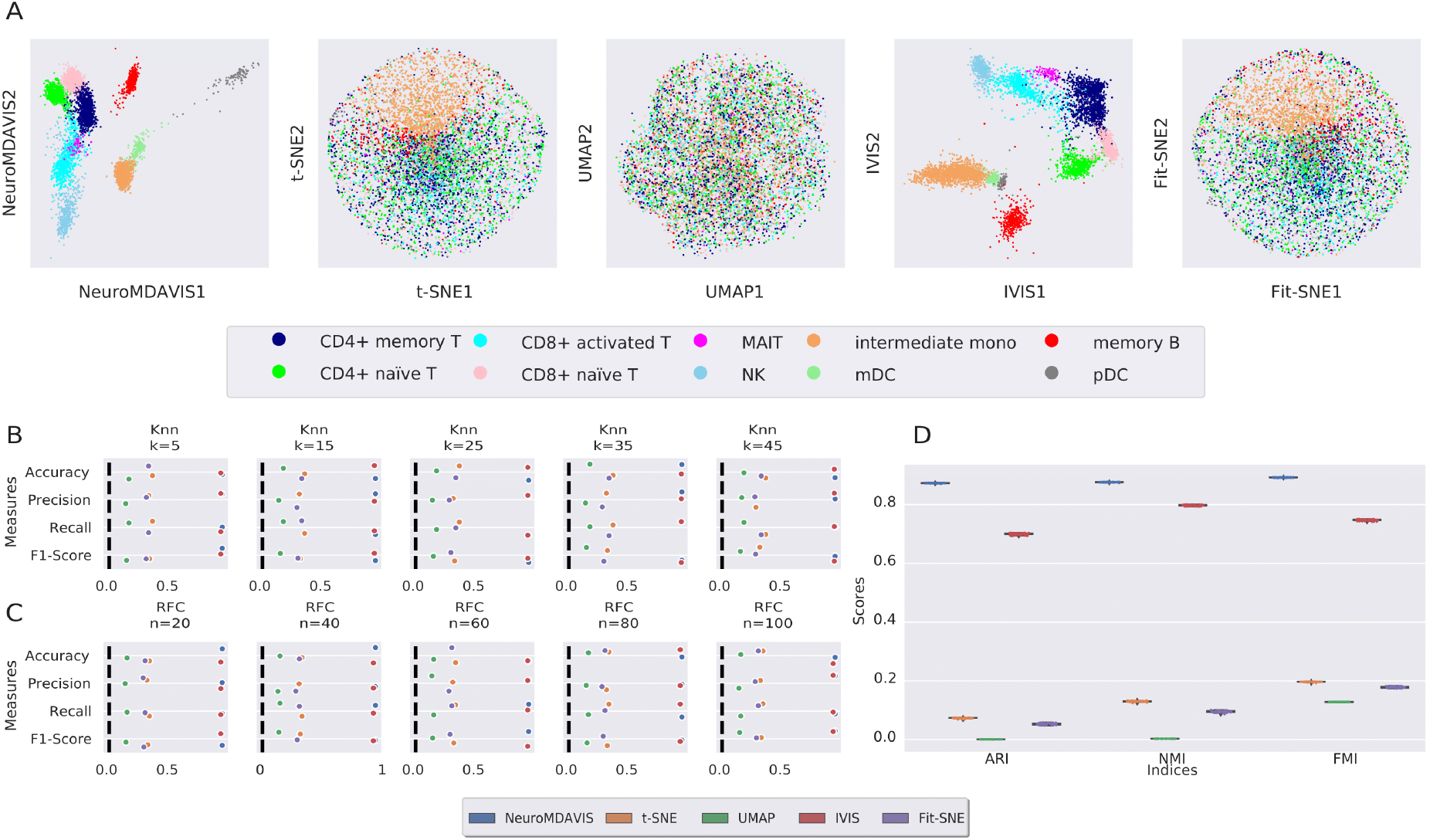
(A) 2-dimensional embeddings produced by NeuroMDAVIS, t-SNE, UMAP, IVIS and Fit-SNE for the multiome *pbmc*10*k* dataset. (B) and (C) show classification performance on projections of *pbmc*10*k* dataset generated by NeuroMDAVIS, t-SNE, UMAP, IVIS and Fit-SNE in terms of accuracy, precision, recall and F1-score using *k*-NN and Random Forest classifers. (D) *k*-means clustering performance on embeddings of *pbmc*10*k* dataset produced by NeuroMDAVIS, t-SNE, UMAP, IVIS and Fit-SNE in terms of ARI, NMI and FMI scores.

Additionally, to evaluate the quality of the embedding quantitatively, we have performed further downstream analysis, i.e., classification and clustering of cell-types on the NeuroMDAVIS-generated embedding. A similar train-test split has been used for classification as followed in case of CITE-seq data. Classification results on the held out test dataset, as shown in Figures 6(B) and 6(C), demonstrate NeuroMDAVIS and IVIS as the best performing models for dimension reduction and visualization across all measures like accuracy, precision, recall and F1-score. On a similar note, *k*-means clustering on the NeuroMDAVIS projection has produced the highest ARI, FMI and NMI scores compared to all other state-of-the-art methods, like t-SNE, UMAP, IVIS and Fit-SNE, as shown in Figure 6(D).

### 3.4 Projection of new observations

Parametric models are always one step ahead when it comes to visualization since they can be used as pre-trained models. NeuroMDAVIS, being a parametric model, can also be used as a pre-trained model to visualize data that are not present during the training process. In order to evaluate NeuroMDAVIS for its capability to visualize new observations not present during training, we have used both CITE-seq and multiome datasets. For each of these datasets, following a 60:40 split of training:test datasets, NeuroMDAVIS has been trained on the 60% training data and the held-out test data of 40% samples has been used for visualization of new observations. As shown in Figure 7(A), for the *bmcite*30*k* dataset, the embeddings on training samples and on test samples are very similar. In both the embeddings, *CD*8, *CD*4, *CD*14, *NK* and *B* celltypes are clearly visible. Similar results have been observed on *kotliarov*50*k* dataset too (Figure 7(B)). All the cell-types are identifiable in the both the training and test embeddings, with their distributions being quite close to each other.

**Figure 7:**
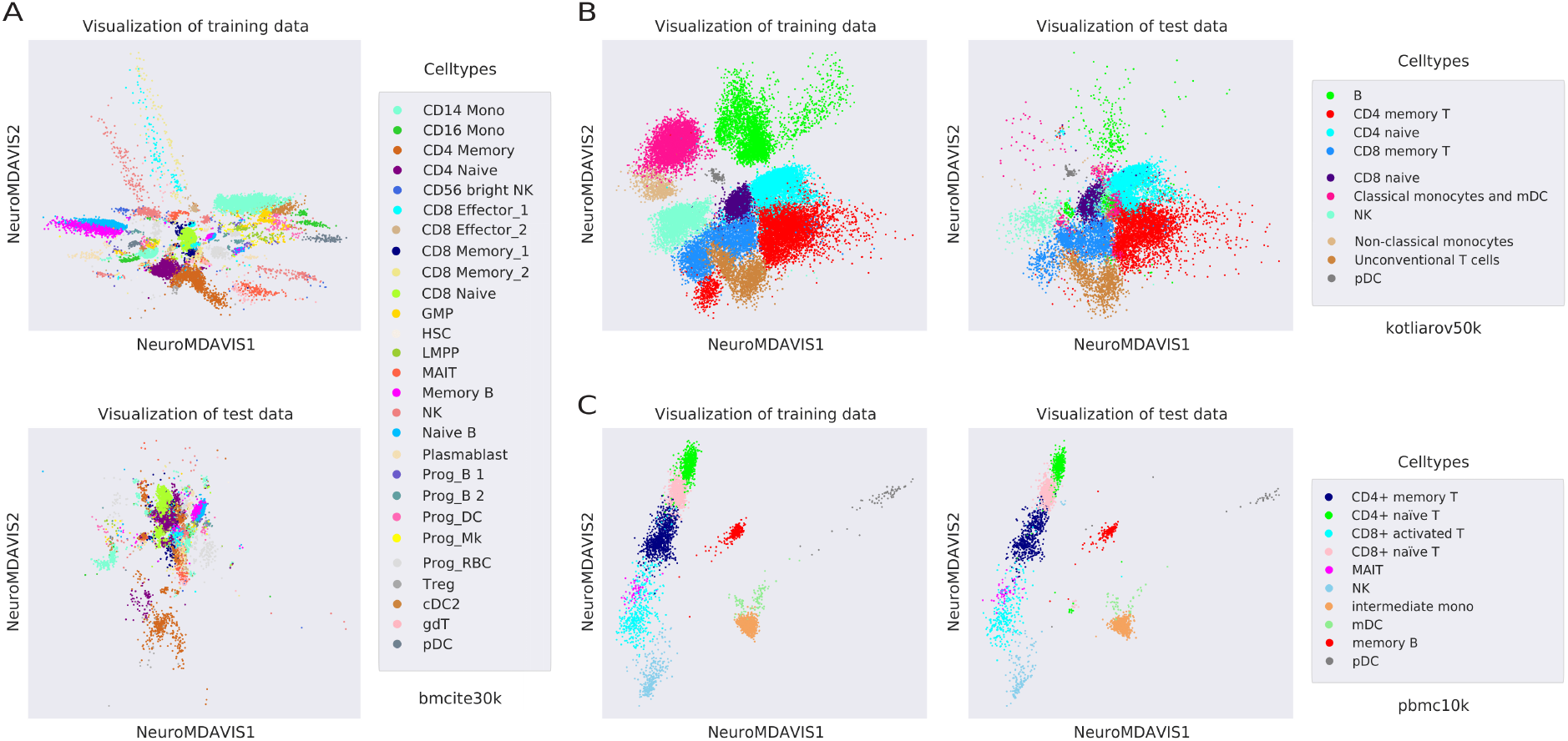
(A), (B) and (C) show 2D embeddings produced by NeuroMDAVIS on both the training and test datasets for *bmcite*30*k, kotliarov*50*k* and *pbmc*10*k* respectively.

For the multiome dataset, as depicted in Figure 7(C), the major cell types, viz., *T, NK* and *B*, are clearly identifiable in both the training and test embeddings. All sub-types of *T* cells are also well separated. Thus, even when the entire data is not available at the time of training, NeuroMDAVIS supports visualization of newer observations added to an existing embedding at runtime.

## 4 Discussion

In this work, we have developed a neural network model, called NeuroMDAVIS, for multi-omics data visualization. NeuroMDAVIS provides joint visualization combining all the omics modalities, and is the first of its kind in this regard. The model is a generalization of NeuroDAVIS [23] recently developed by the authors for the purpose of visualizing single data modality. The performance of NeuroMDAVIS has been demonstrated on CITE-seq and multiome datasets and results have been compared with several state-of-the-art methods, like t-SNE, UMAP, IVIS and Fit-SNE. Additionally, we have also benchmarked NeuroDAVIS on multiple biological datasets with single modality, like RNA-seq and Mass Cytometry data. Both NeuroDAVIS and NeuroMDAVIS are non-recurrent, feed forward neural network architectures that can be used to produce a latent 2-dimensional embedding from high-dimensional single-omics and multi-omics data respectively. This latent embedding/projection captures significant data characteristics which can be useful for various downstream tasks including classification and clustering.

For single-cell omics data with single modality, NeuroDAVIS has shown better local shape preservation capability than the other existing methodologies. Different subsampling-based experiments have been carried out to test robustness of NeuroDAVIS, which it has passed successfully. Moreover, the capability of local neighbourhood preservation in projected space of NeuroDAVIS has been found to be the best in terms of trustworthiness (a measure of local neighbourhood preservation), among all existing methods used for visualization. NeuroDAVIS has also been able to represent the differentiation states across cells for marker proteins in the Mass Cytometry data. In the context to multi-omics data visualization, NeuroMDAVIS has been able to produce qualitatively excellent projections compared to the other state-of-the-art visualization methods. The latent embedding when used for cell-type classification or clustering has surpassed those obtained by most of the other methods.

Overall, both NeuroDAVIS and NeuroMDAVIS have shown excellent dimension reduction and visualization capability on biological datasets, most importantly, without the usage of PCA as a prior initialization step, unlike other existing methods. One shortcoming of both these models is that they take an identity matrix whose order is equal to the number of samples, as input, thereby consuming large memory space. However, both these models are parametric and do not assume any prior data distribution, which stands out as one of their key strengths. Further, to our knowledge, there has been no single method till date that supports joint visualization of multi-omics data. Researchers have been visualizing one omics layer overlaid on the other. NeuroMDAVIS aims to fill up this gap. Moreover, unlike major existing visualization methods like t-SNE, UMAP, both NeuroDAVIS and NeuroMDAVIS can be used as pretrained models to visualize streaming samples, which implies that they support iterative continuous integration and deployment. This attribute makes them highly suitable for projects like Human Cell Atlas (HCA) or Human Tumor Atlas network (HTAN) initiatives. Thus, NeuroDAVIS and NeuroMDAVIS together can be claimed to be the new state-of-the-art for omics data visualization.

## Acknowledgements

This work is supported by DST-NSF grant provided to RKD through IDEAS-TIH, Indian Statistical Institute, Kolkata, India.

## Authors’ contributions statement

Conceptualization of methodology and framework: CM, DBS, VD, RKD. Data curation, Data analysis, Formal analysis, CM, DBS, VD. Implementation: CM. Investigation, Code review, Validation, Initial draft preparation: CM, DBS. Reviewing, Editing, Overall Supervision: VD, RKD.

## Conflict of Interest

VD works as a Lead Data Scientist at Novo Nordisk A/S, Måløv. DBS works as a postdoctoral researcher at German Cancer Research Center (DKFZ), Heidelberg, Germany. They have not received any funds for this work.

## Data and Code availability

The datasets used in this study can be downloaded from https://doi.org/10.5281/zenodo.10623932. Codes to reproduce the results can be found at https://github.com/shallowlearner93/NeuroMDAVIS.

## Notes

https://doi.org/10.5281/zenodo.10623932

https://github.com/shallowlearner93/NeuroMDAVIS

### Competing Interest Statement

VD works as a Lead Data Scientist at Novo Nordisk A/S, Malov. DBS works as a postdoctoral researcher at German Cancer Research Center (DKFZ), Heidelberg, Germany. They have not received any funds for this work.

